# FHL-1 interacts with human RPE cells through the α5β1 integrin and confers protection against oxidative stress

**DOI:** 10.1101/2020.09.28.317263

**Authors:** Rawshan Choudhury, Nadhim Bayatti, Richard Scharff, Ewa Szula, Viranga Tilakaratna, Maja Søberg Udsen, Selina McHarg, Janet A Askari, Martin J Humphries, Paul N Bishop, Simon J Clark

## Abstract

Retinal pigment epithelial (RPE) cells that underlie the neurosensory retina are essential for the maintenance of photoreceptor cells and hence vision. Interactions between the RPE and their basement membrane, *i.e*. the inner layer of Bruch’s membrane, are essential for RPE cell health and function, but the signals induced by Bruch’s membrane engagement, and their contributions to RPE cell fate determination remain poorly defined. Here, we studied the functional role of the soluble complement regulator and component of Bruch’s membrane, Factor H-like protein 1 (FHL-1). Human primary RPE cells adhered to FHL-1 in a manner that was eliminated by either mutagenesis of the integrin-binding RGD motif in FHL-1 or by using competing antibodies directed against the α5 and β1 integrin subunits. The results obtained from primary RPE cells were replicated using the hTERT-RPE cell line. RNAseq expression analysis of hTERT-RPE cells bound to FHL-1 showed an increased expression of the heat-shock protein genes *HSPA6, CRYAB, HSPA1A* and *HSPA1B* when compared to cells bound to fibronectin (FN) or laminin (LA). Pathway analysis implicated changes in EIF2 signalling, the unfolded protein response, and mineralocorticoid receptor signalling as putative pathways. Subsequent cell survival assays using H_2_O_2_ to induce oxidative stress-induced cell death showed hTERT-RPE cells had significantly greater protection when bound to FHL-1 or LA compared to plastic or FN. These data show a non-canonical role of FHL-1 in protecting RPE cells against oxidative stress and identifies a novel interaction that has implications for ocular diseases such as age-related macular degeneration.

## Introduction

The retinal pigment epithelium (RPE), a monolayer of cells in the retina, makes an essential contribution to the maintenance and support of the photoreceptor cells, and hence vision itself (1). Disruption to the normal homeostasis of RPE cells is linked to a range of retinal degenerative diseases including age-related macular degeneration (AMD) (2): the third leading cause of blindness in the world (3). RPE cells have a high metabolic turnover and are exposed to extreme levels of light-induced oxidative stress (4). Furthermore, these cells are among the most actively phagocytic cells in the body where they deal with the constant shedding of the outer segments of both rod and cone cells, which are dynamic structures that undergo constant renewal (5). RPE cells also play fundamental roles in the transportation of nutrients from the underlying blood vasculature (termed the choriocapillaris, see Figure 1a) to photoreceptors, and *vice versa*, such as the transportation of ions, water and metabolic end-products from the sub-retinal space to the blood.

**Figure 1.**
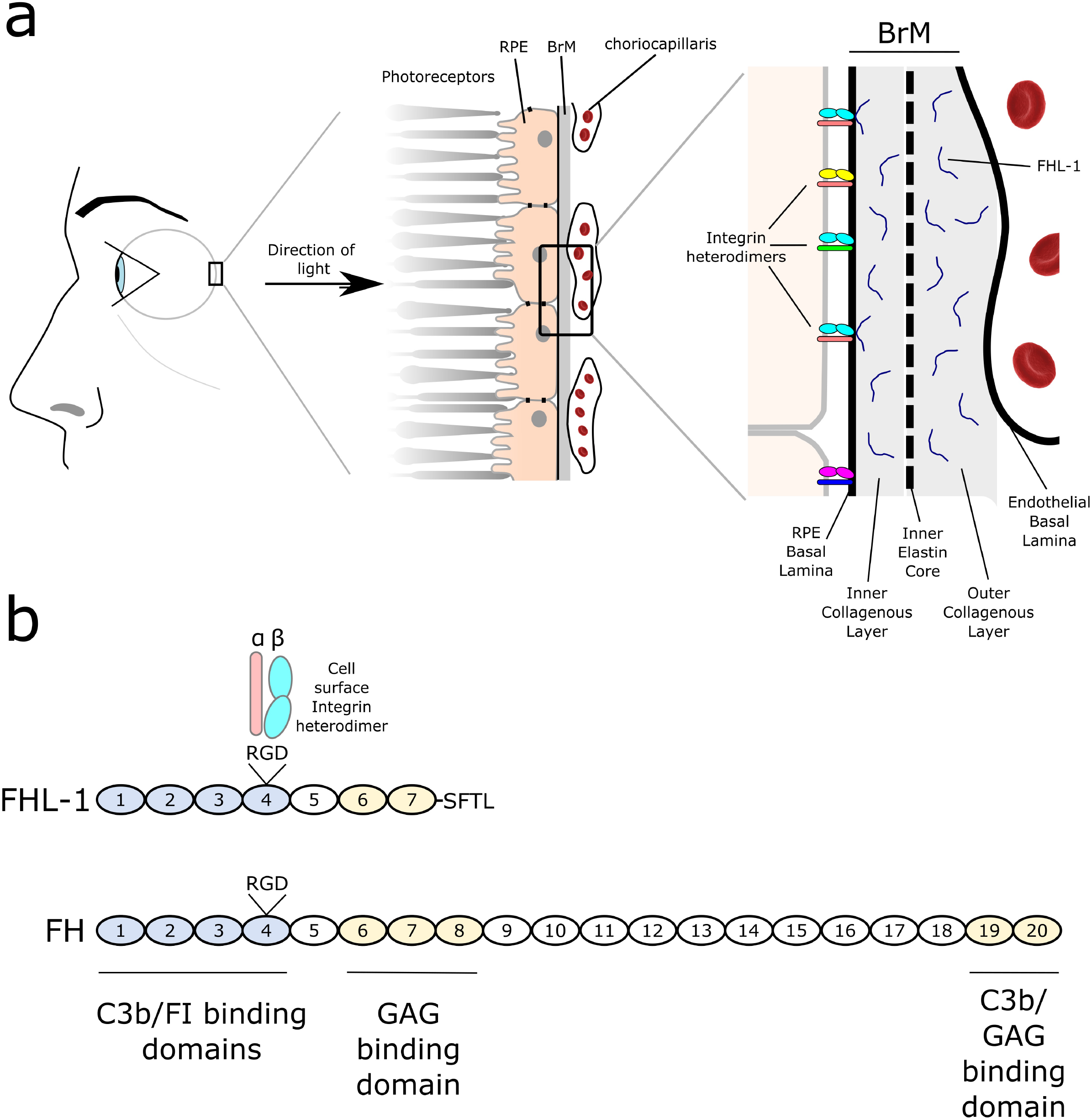
Schematic of RPE cell interactions with their underlying ECM. **a**) The RPE forms a monolayer on Bruch’s membrane and plays a crucial role in maintaining photoreceptors. Bruch’s membrane separates the RPE its blood supply, the choriocapillaris. RPE cells adhere to, and interact with, ligands within the basement membrane of Bruch’s membrane (BrM), including FN and LA, through various integrin heterodimer receptors expressed on their surface. Bruch’s membrane comprises five separate layers: the RPE basement membrane; the inner collagenous layer; an elastin core; the outer collagenous layer; and the endothelial basement membrane layer. The complement regulator FHL-1 is found anchored to heparan sulphate and dermatan sulphate glycosaminoglycan (GAG) chains found in collagenous and basement membrane layers of Bruch’s membrane. **b**) Both FH and its truncated variant FHL-1 are comprised of complement control protein (CCP) domains: FH has twenty domains, while FHL-1 has seven CCPs identical to FH but with a unique C-terminal tail. Both proteins also share an RGD integrin-binding motif in CCP4.

RPE cells are separated from the choriocapillaris by an acellular barrier called Bruch’s membrane (Figure 1a). This extracellular matrix (ECM) is comprised of five separate layers; the RPE basement membrane, inner collagenous layer, an elastin core, the outer collagenous layer, and the choriocapillaris basement membrane (6). The structure and permeability of Bruch’s membrane is important for maintaining a healthy environment in the eye. Bruch’s membrane itself confers some selectivity as to what can, or cannot, pass through from the choroid to the retinal space (7), leading to two immunologically semi-independent regions. Indeed, the breakdown of Bruch’s membrane barrier function allows the intrusion of blood vessels into the retinal space and is the site of lipid and debris accumulation that leads to the formation of drusen, the hallmark lesions of the early stages of AMD (8).

The attachment of RPE cells to their underlying Bruch’s membrane is required to maintain their homeostasis, and changes to this ECM affects RPE cell gene transcription and protein secretion (9). Attachment of the RPE to the inner basement membrane of Bruch’s membrane is mediated at least in part by integrins (Figure 1a). Integrins are heterodimeric proteins comprising various combinations of α and β subunits that determine ligand-binding specificity (10, 11) and control the morphology and fate of the host cell (12). A variety of integrins are expressed by the RPE (13), including the α5β1 integrin (*ITGA5:ITGB1*) which has been shown to mediate cell attachment, migration and proliferation (14). The α5β1 integrin recognises the conserved binding motif Arg-Gly-Asp (RGD) that is present in various ECM ligands including fibronectin (FN) (15).

Complement factor H-like protein 1 (FHL-1) is a truncated form of the complement inhibitory protein factor H (FH), which arises from alternative splicing of the *CFH* gene on chromosome 1 (16, 17). FHL-1, and to a lesser extent FH, have been identified within Bruch’s membrane and at the interface between Bruch’s membrane and the RPE (7, 18). Genetic variants in the *CFH* gene are associated with increased risk of AMD (19, 20) and this is thought to be due to decreased activity of FHL-1 and FH that results in increased complement activation in the ECM, leading to a local inflammatory response, the recruitment of circulating immune cells, formation of drusen, and ultimately RPE cell death (21–23). However, recent studies have begun to identify non-canonical roles of FH and FHL-1 and the potential contribution of their dysregulation to AMD pathogenesis through inducing RPE mitochondrial dysfunction (24, 25) and lipid peroxidation (26). Both FH and FHL-1 have an RGD motif on the surface of their fourth complement control protein (CCP) domain (see Figure 1b) and it has previously been demonstrated that FHL-1 can confer cell attachment activity to human epithelial and fibroblast cell lines *via* this RGD motif (27).

Herein, we investigate the interactions between primary human RPE cells, and the immortalised RPE cell line hTERT-RPE1, with immobilised FHL-1. We investigate the role of RPE cell integrins in interacting with FHL-1 and, by using RNAseq and cell survival assays, analyse the downstream consequences of the RPE cell/FHL-1 interaction compared to basement membrane components including laminin (LA) and FN. Subsequently, we elucidate a novel role for RPE/FHL-1 interactions in RPE cell modulation and their survival in response to oxidative stress.

## Materials and Methods

### Primary RPE isolation from human eye-globes

Human eyes were collected from the Manchester Royal Eye Hospital Eye Bank after removal of the corneas for transplantation. Consent had been obtained for the eye tissue to be used for research and guidelines established in the Human Tissue Act of 2004 (UK) were adhered to. Ethical approval for the use of human donor eyes was given by North West – Greater Manchester Central Research Ethics Committee (REC reference 15/NW/0932). A total of 22 donor eye pairs were used (13 female, 9 male) with an age-range between 42 and 77 years old (mean age 58.8 years old). All donors used were <48 hours post mortem. Eye globes were collected in collection media (Hank’s Balanced Salt Solution (Sigma-Aldrich, Poole, UK) supplemented with 1% (w/v) Amphotericin-B, 1% (w/v) Penicillin/Streptomycin, 0.5% (w/v) Sodium Pyruvate, 1% (w/v) HEPES, 0.01 mg/ml gentamycin (Sigma-Aldrich). Each eye globe was placed in a 10 cm petri dish and washed with phosphate-buffered saline (PBS). Then the iris, lens, vitreous and neurosensory retina (NSR) were gently removed from the posterior eye-cup using a scalpel, forceps and scissors. After that, the eye cup was rinsed with sterile PBS and the interior was digested using 0.5 mg/ml each of collagenase Type I-A (from *Clostridium histolyticum*, Sigma-Aldrich) and Type IV (from *Clostridium*, Sigma-Aldrich) in 5 ml of sterile-filtered Dulbecco’s Modified Eagle’s Medium (DMEM3, Sigma-Aldrich) media with high glucose for 90 min at 37°C. Next the media containing the collagenases in the eye cup was gently discarded, then the eye cup was filled with 5 ml of fresh DMEM media and the RPE were gently scraped off the underlying Bruch’s membrane.

The media containing RPE sheets/fragments was collected and centrifuged at 150 g for 5 min. The supernatant was removed and the RPE sheets/fragments were resuspended in 4 ml RPE media (1:1, DMEM3: Ham’s F12 medium (GIBCO 31330-038), 10% (v/v) foetal calf serum, 1% (w/v) penicillin-streptomycin). Resuspended RPE was added to 0.5% (w/v) gelatin (Sigma-Aldrich)-PBS coated-wells of 12 or 24 well plates. After 24h, the RPE was washed 3x with PBS and fresh media was added. RPE cells were grown until they were confluent and ready to be passaged.

### Clonal cell line tissue culture

The human immortalized RPE cell line hTERT-RPE (hTERT-RPE-1, ATCC®CRL-400) was purchased from ATCC (LGC Standards, UK). The hTERT-RPE-1 cells were grown in DMEM: Ham’s F-12 (ATCC, 1:1) media containing 10% (v/v) foetal calf serum (ATCC), 100 U/ml penicillin and 100 mg/ml streptomycin and 0.01 mg/ml hygromycin B (Sigma-Aldrich). HEK293T cells were grown in DMEM3, containing 10% (v/v) fetal calf serum, 100 U/ml penicillin and 100 mg/ml streptomycin. All cells were maintained at 37°C in 5% humidified CO_2_.

### Transfection of wild-type and RGD-null FHL-1 plasmid into HEK293T

cDNA encoding histidine-tagged full-length wild-type FHL-1 (18) and the RGD-null FHL-1 protein, where the aspartic acid is substituted with an alanine to disrupt the RGD binding site was synthesised commercially (Life Technologies, Paisley, UK). Purified cDNA was incorporated into a pcDNA3.1 vector and transfected into HEK293T cells using PEI max transfection reagent (Polysciences, Germany) as described previously (18). Briefly, for each 15 cm dish, 1 mg/ml plasmid DNA and 7.5 mM PEI were added to separate aliquots of 150 mM NaCl and each incubated for 10 min at room temperature. After incubation, the PEI solution was slowly added to the solution containing DNA and then incubated for 10 min at room temperature. The DNA/PEI mixture was added to a 15 cm diameter dish containing 7×10^6^ HEK293T cells/per dish in a dropwise manner and incubated at 37°C. After 5-6 hours, the media was replaced with 2% (v/v) FCS containing DMEM3 (no antibiotic) and incubated at 37°C for overnight. Conditioned media was collected daily and replaced with fresh media over the next 4 days.

### Purification of wild-type and RGD-null FHL-1

Conditioned media from each of the 20 dishes (17.5ml/dish) were collected and pooled after 24, 48, 72, and 144 hours. Total conditioned media after 144 hours (1,400 mls total) was diluted with the addition of 600 mls 50 mM HEPES, 500 mM NaCl, 20 mM Imidazole, pH 7.5. To this, 12 ml of NiNTA resin (Expedeon) was added and incubated overnight with rotation at 4°C. The NiNTA beads were collected by passing the media through empty PD10 columns with a filter (GE Healthcare) by gravity flow (500 ml media per PD10 column, i.e. four PD10 columns in total). The beads were then washed with 10 ml of wash buffer (50 mM HEPES, 500 mM NaCl, 20 mM Imidazole, pH 7.5). Finally, wild-type FHL-1 or RGD-null FHL-1 protein was eluted using 16 ml of elution buffer (50 mM HEPES, 500 mM NaCl, 500 mM imidazole, pH 7.5). Eluted protein was dialysed back into wash buffer overnight at 4°C before being concentrated further by addition to 1.5ml NiNTA beads and eluted in 6 x 1ml aliquots. Purified protein aliquots were dialysed into 20 mM glycine, 125 mM NaCl, pH 9.0 using a dialysis cassette Slide-A-Lyzer (Fisher, cat no 10759784) with a 10 kDa cut off. The purified recombinant proteins (i.e. FHL-1 and RGD-null FHL-1) were assessed for purity by SDS-PAGE, and visualised by staining the gels for 60min at room temperature with Instant Blue Coomassie stain (Expedeon, Cambridge, UK).

### Endotoxin testing of purified FHL-1 proteins

The endotoxin level of FHL-1 recombinant protein preparations was measured using the Toxin Sensor™ Chromogenic LAL Endotoxin Assay Kit (GenScript, NJ, USA), according to the manufacturer’s protocol. This method utilizes a modified Limulus Amebocyte Lysate (LAL) and a synthetic colour producing substrate to detect endotoxin chromogenically. Briefly, 100 μl of standards (0, 0.01, 0.025, 0.05, 0.1 and 0.5 EU/ml), test samples (recombinant wild-type FHL-1 and RGD-null FHL-1 proteins) and a blank containing 100 μl of LAL reagent water were dispensed into endotoxin-free vials in duplicate. 100 μl of reconstituted LAL was added to each vial, mixed by swirling and incubated at 37°C in a water bath. After incubation, 100 μl of reconstituted chromogenic substrate solution was added to each vial and mixed gently to avoid foaming. All the vials were further incubated at 37°C for 6 minutes. Finally, 500 μl of each of three colour stabilisers were added eventually to each vial and mixed gently to stop the reaction. The absorbance of each reaction vial was read at 545 nm.

Under the standard conditions, the absorbance at 545 nm shows a linear relationship within a concentration in the range of 0.01 to 1 EU/ml. The absorbance for the five standards was plotted on the x-axis and the corresponding endotoxin concentration in EU/ml on the y-axis. A best-fit straight line was drawn between these points and the endotoxin concentrations of samples were determined graphically.

### Fluid-phase cofactor activity of wild-type and RGD-null FHL-1 (C3b break-down assay)

To test the functional capacity of both FHL-1 and RGD-null FHL-1, a C3b breakdown assay was employed as described previously (23). Briefly, 0.1 μg of either wild-type FHL-1 or RGD-null FHL-1 was incubated with 2 μg C3b (VWR International, Lutterworth, UK), and 0.4 μg factor I (FI) (VWR International,) in PBS (total volume of 20 μl) for 15 minutes at 37°C. The reaction was stopped by the addition of 5 μl 5x SDS reducing sample buffer (NuPAGE LDS sample buffer, Life Technologies) and boiling for 10 minutes at 100°C. Samples were run on a 4-12% NuPAGE Bis Tris gels (Life Technologies, UK) at 150V for 75 minutes to allow the separation of the C3b breakdown product bands. Blue Protein Standard Broad Range (New England Biolabs, Hitchin, UK) was used as a protein marker. The gels were stained using Instant Blue for 1 hour at room temperature. Gel images were taken using an Alpha Innotech FluorChem 5500 gel imaging system.

### Immunofluorescence staining

Primary RPE cells were seeded on coverslips and grown until they were confluent. Cells were fixed in 4% (w/v) paraformaldehyde (PFA) for 20 min at 4°C. PFA was removed and the coverslips washed with PBS. Cells were permeabilised with 0.5% (v/v) Triton X-100 in PBS for 10 min at room temperature. After washing with PBS, the seeded coverslips were blocked with 5% (v/v) normal goat serum (NGS) for 1h at room temperature. These were then washed three times with PBS and labelled with antibodies directed against RPE cell markers including mouse monoclonal anti-RPE65 (Abcam, clone: 401.8B11.3D9) and anti-bestrophin-1 (Novus Biologicals, UK, clone: IgG1 E6-6), or the tight-junction marker anti-ZO-1 (Invitrogen, clone: ZO-1-IA1Z). All antibodies were diluted 1:50 in PBS containing 5% (v/v) normal goat serum (NGS; 100 μl/coverslip) (Novus Biologicals, UK) and incubated overnight at 4°C. Cells were washed with PBS and then incubated with secondary antibody Alexa Fluor 488-conjugated goat anti-mouse antibody (Life Technologies) diluted at 1:100 in 5% (v/v) NGS in PBS for 1 hour at room temperature. Finally, DAPI was applied as a nuclear counter stain at 0.3 μM for 5 minutes prior to mounting with medium (Vectashield soft, Vector Labs, Peterborough, UK) and placing the coverslips upside down onto microscope slides. Images were taken using by a snapshot widefield microscope (Leica, 20x/0.50 PL FLUVOTAR objective) using HCImage software.

### Cell spreading assay

For the cell spreading assays, 96-well plates were coated with different protein ligands (50 to 80 μg of FHL-1, FH (HyCult, Uden, The Netherlands), FHR-4 (23) and 10 μg of FN (Sigma-Aldrich, cat F1141) and BSA) in PBS (with Ca^2+^ and Mg^2+^, Sigma-Aldrich) and incubated overnight at 4°C. Then all of the coated wells were blocked with heat denatured 1% BSA in PBS (heated for 11 min at 85°C to denature the BSA) for one hour at room temperature. Primary RPE or hTERT-RPE-1 cells were trypsinized, counted using haemocytometer and 10,000 cells per well added for each condition in duplicate or in triplicate, prior to incubating for 3 hours at 37°C. The cells were photographed every hour (Leica microscope, 10x/0.22 HI PLAN objective and DFC420 camera) with at least four images being taken for each well; these images were analysed using Leica imaging software and ImageJ to assess the degree of spreading. Cells were defined as spread if they were phase dark and exhibited visible cytoplasm all around the nucleus.

### Inhibition and Competition assay

96-well plates were coated with different protein ligands (80 μg FHL-1, FH, FHR-4 and 10 μg of FN and BSA) in PBS (with Ca^2+^ and Mg^2+^, Sigma-Aldrich) and incubated overnight at 4°C. Then all coated wells were blocked with heat denatured 1% (w/v) BSA in PBS for one hour at room temperature. Primary RPE and hTERT-RPE-1 cells were trypsinized and counted using a haemocytometer. Cells (in each condition 10,000 cells/well in triplicate) were incubated with 10 μg/ml of mouse monoclonal anti-α5 (mAb16), anti-β1 (mAb 13), anti-αV (17e6), anti-αVβ3 (LM609) or anti-αVβ5 (PIF6) integrin antibodies (all kindly supplied by the Prof. Humphries lab), for 20 min at room temperature before adding to the wells and incubated for 3 hours at 37°C.

In other inhibition assays, cells were incubated with increasing concentrations (0 to 120 μg/ml) of a reverse sequence control peptide (Ser-Asp-Gly-Arg-Gly/SDGRG, Sigma-Aldrich) and an RGD peptide (Gly-Arg-Gly-Asp-Ser/GRGDS, Sigma-Aldrich) for 20 min at room temperature before the cells were added to the wells for incubation for 3 hours at 37°C.

In competition assays, either primary RPE or immortalized hTERT-RPE-1 cells were pre-incubated with different proteins such as FHL-1 (50 μg/ml), CCP6-7 of FHL-1 (28) at 50 μg/ml, FH (150 μg/ml) and FN (10 μg/ml) for 20 min at room temperature before being added to FHL-1 (50 μg/ml) coated plates. Cells were incubated at 37°C for up to 3 hours.

### RNA isolations and quantitative RT-PCR (qPCR)

The hTERT-RPE-1 cells (1x 10^5^ cells per well) were grown on either 50 μg/ml FHL-1, 10 μg/ml FN or 10 μg/ml LA (Merck Millipore, cat. No. CC095) substrate in a 24-well plate for 24 hours at 37°C. For gene expression analysis, total RNA was extracted from the attached cells using an RNA Isolate II Mini Kit (BioLine) according to the manufacturer’s protocol. RNA quality and concentration were measured with a NanoDrop 2000 spectrophotometer (ThermoFisher Scientific). cDNA was synthesized using 1 μg of RNA and a High Capacity cDNA Reverse Transcription kit (Applied Biosystems) according to the manufacturer’s instructions. PCR was carried out using TaqMan®Gene Expression Assays (FAM/MGB-NFQ, ThermoFisher Scientific, Paisley, UK) and real time PCR (qPCR StepOne Plus, ThermoFisher) cyclers. PCR reactions were carried out on 96-well plates (MicroAmp®Fast Optical 96-well Reaction plate, Applied Biosystems, Life Technologies) and 2 μl cDNA (10-20 ng diluted in nuclease-free water) was added to each well in triplicate.

### Gene Transcriptomic analysis by RNA-seq

For RNA-seq, hTRERT-RPE1 cells were grown to 80% confluency on either FHL1 (20μg/ml), FN or LA (10μg/ml). For each analysis 2 wells of a six-well plate were used and 3 replicates were performed (n = 9 in total). After media was fully aspirated from the wells, total RNA was isolated using an Isolate II RNA Mini Kit (Bioline) directly from the wells as per manufacturer’s instructions, and RNA quality and concentration measured using a Nanodrop 2000 spectrophotometer. Unmapped paired-end sequences from an Illumina HiSeq4000 sequencer were tested by FastQC (http://www.bioinformatics.babraham.ac.uk/projects/fastqc/). Sequence adapters were removed and reads were quality trimmed using Trimmomatic_0.36 (29). The reads were mapped against the reference human genome (hg38) and counts per gene were calculated using annotation from GENCODE 27 (http://www.gencodegenes.org/) using STAR_2.5.3 (30). Normalisation, Principal Component Analysis, and differential expression were calculated with DESeq2_1.16.1 (31). Differential transcriptional analysis was carried out using Ingenuity Pathway Analysis (Qiagen) comparing differences between substrates in a core analysis with a global false discovery rate (FDR) of 0.1. Analysis was also carried out using DAVID (https://david.ncifcrf.gov/) with a fold change cut-off of 1.5 when comparing between gene expression between substrates, transcripts with counts <50 were defined as being absent.

### Cell Survival Assay

hTERT-RPE1 cells were maintained in DMEM:F12 (1:1) media supplemented with 10% (v/v) FCS, 1% (w/v) penicillin-streptomycin and 0.01 mg/ml hygromycin B (all from Thermo Fisher Scientific UK). For experiments, cells were plated at a density of 100,000cm^-2^ in 96 well plates (Corning, MERCK UK), with wells previously coated for 24h with 20μg/ml FHL-1, 20μg/ml RGD null FHL-1, 10μg/ml FN (Sigma-Aldrich), 10μg/ml LA (Sigma-Aldrich) or 1x PBS with added Ca^2+^ and Mg^2+^ (ThermoFisher Scientific UK) vehicle. After a further 24h cells were switched to serum-free DMEM:F12 (1:1) medium supplemented with B27 without antioxidants (Thermo Fisher Scientific, UK). Cells from each group of coated wells were incubated in the above media in the presence or absence of 150μM H_2_O_2_ (Sigma-Aldrich) for a further 24h. Cells were then fixed in 4% paraformaldehyde (Sigma-Aldrich) in 1x PBS for 15mins at RT and washed three times in PBS. During the second wash, 1μg/ml Hoechst 33342 (Sigma-Aldrich) added to the PBS wash to stain nuclei. Cells were imaged using a CellInsight CX5 scanner (Thermo Fisher Scientific) with a 10x objective, taking three snapshots per well in identical positions, and three wells for each condition per experiment for three experiments. Cell counting was carried using the ImageJ software package. After thresholding to create a binary image, cell nuclei sized between 0-150 pixel^^2^ were included in counts. Numbers using these settings were consistent with manual counting from images from PBS control and PBS H_2_O_2_-stimulated cells.

## Results

### Primary human RPE cells interact with immobilised FHL-1

To investigate potential interactions between RPE cells and FHL-1, primary RPE cells were isolated from human donor cadaver eyes. Cells were examined for RPE marker expression by immunofluorescence, including ZO-1, RPE-65 and bestrophin-1 (Supplementary Figure 1). Cell spreading assays were used to determine whether the primary RPE cells interact with FHL-1 immobilised onto a plastic surface. RPE cells were added to plates pre-treated with immobilised full-length FH or FHL-1 together with a FN positive control, and BSA and FHR-4 (a structurally related protein, but with no native RGD integrin binding motif) as negative controls. After three hours incubation, 40% primary RPE cell spreading was observed on FHL-1 compared to the FN control (Figure 2). In contrast, primary RPE cells did not spread on immobilised full-length FH, despite it sharing an identical RGD binding site with FHL-1. Furthermore, no primary RPE cell spreading was visible on either plastic alone, BSA or FHR-4 (Figure 2). To investigate further the lack of interaction with FH, cell spreading experiments were repeated in the presence of FH as a fluid phase competitor (see Supplementary Figure 2). Additionally, FHL-1, FN and a recombinant protein comprising solely CCPs6-7 of FH were also used as fluid phase competitors. Fluid-phase FH inhibited the spreading of primary RPE cells on immobilised FHL-1, as well as fluid phase FHL-1 and FN (Supplementary Figure 2). This suggests that the previously observed lack of RPE cell interaction with immobilised FH was due to the way the protein was absorbed onto plastic, and subsequent inaccessibility of its RGD domain, rather than an inherent lack of functional activity.

**Figure 2.**
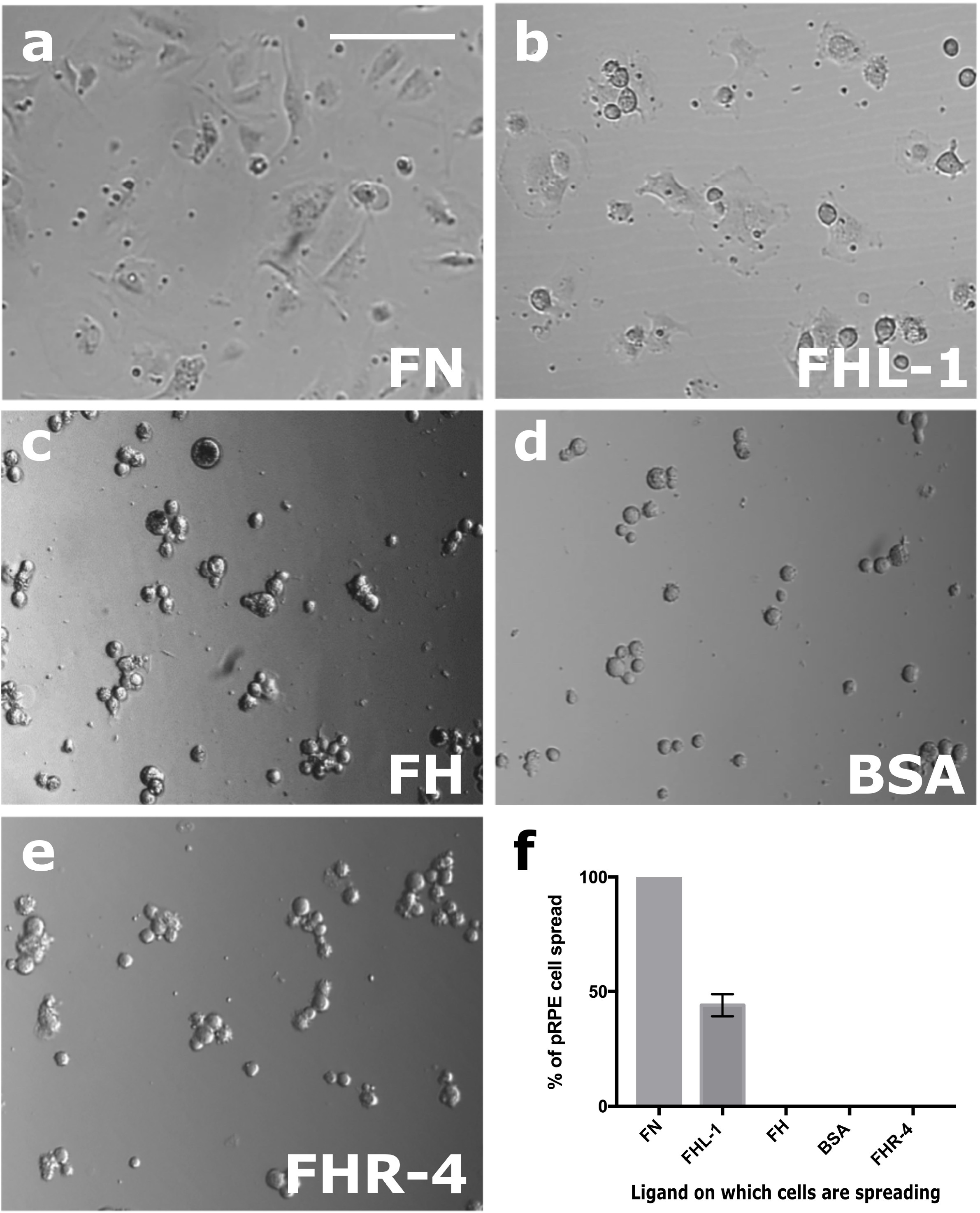
Primary RPE cells interact with immobilised FHL-1. Cultured primary RPE cells collected from human donor eyes were incubated for three hours in wells of 96 well plates coated with: **a**) FN; **b**) FHL-1; **c**) FH; **d**) BSA; or **e**) FHR-4. **f**) the number of spread cells were counted in four different visual fields for each condition and each condition was measured in triplicate. The percentage of spread cells were calculated and compared to the positive control, fibronectin. Scale bar represents 100μm. Images in **a**-**e** are representative of 10 independent experiments. Data in **f** represent n=10 ± s.e.m.

### RPE cell α5β1 integrin interacts with FHL-1 via its RGD site

Given the presence of an RGD motif in CCP4 of FHL-1 (Figure 1b), we tested the role of integrins in FHL-1 binding. Competition assays were performed where primary RPE cells were bound to immobilised FHL-1 in the presence of inhibitory antibodies against integrins known to bind the RGD sequence including anti-α5 (mAb16), anti-β1 (mAb13), anti-αV (17e6), anti-αVβ3 and anti-αVβ5 (Figure 3). The spreading of primary RPE cells on immobilised FHL-1 was completely abolished in the presence of either anti-α5 or anti-β1 antibodies in a dose-dependent manner (Figure 3g-i). In addition, a significant reduction in RPE cell spreading was also achieved with the anti-αVβ5 antibody too, suggesting dual receptor recognition (Figure 3g).

**Figure 3.**
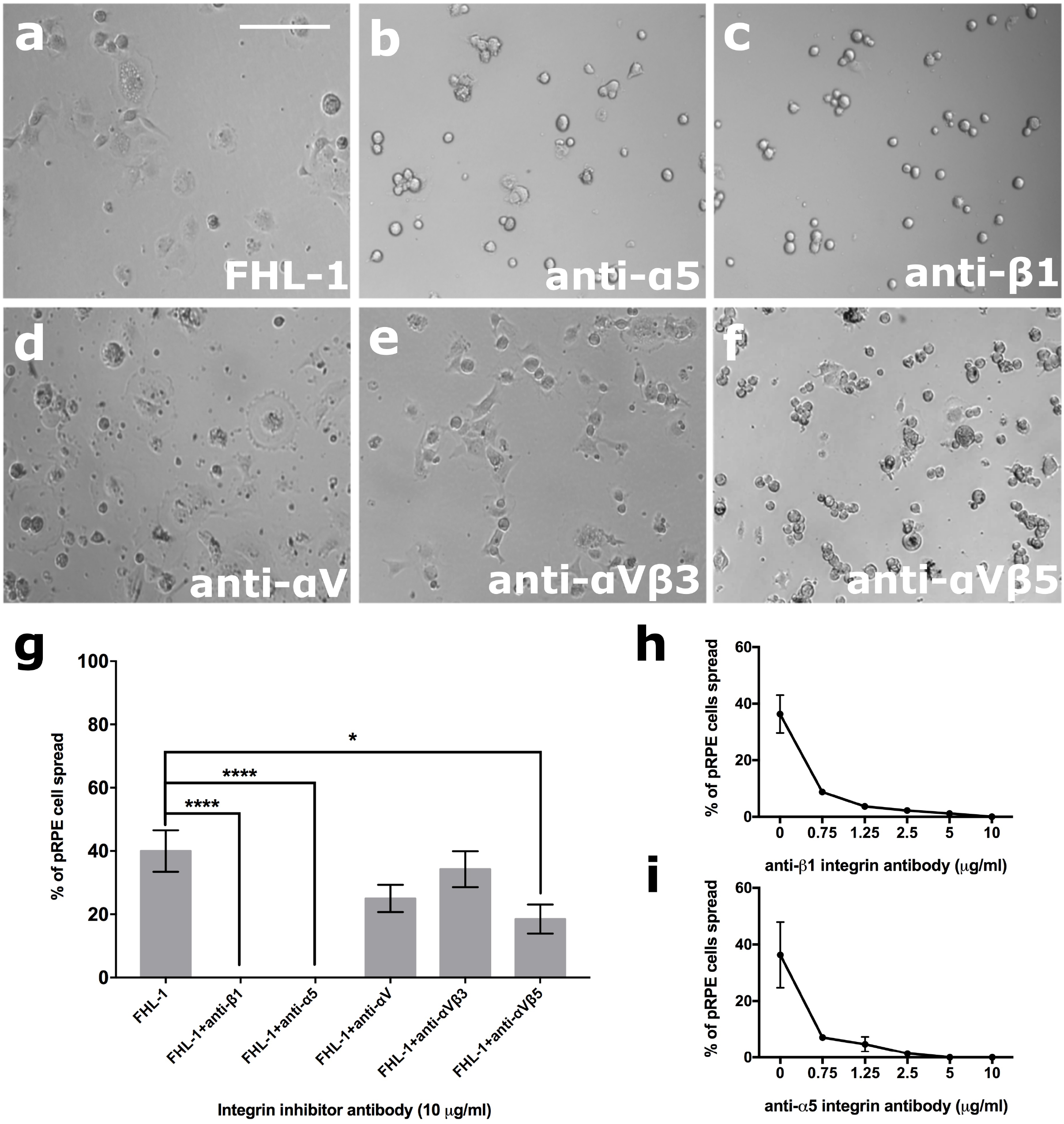
RPE cell interactions with FHL-1 are mediated through the α5β1 integrin. Cultured primary RPE cells were incubated on an FHL-1 matrix (**a**) in the presence of competing antibodies against specific integrin subunits (all 10μg/ml), including: **b**) anti-α5; **c**) anti-β1; **d**) anti-αV; **e**) anti-αVβ3; and **f**) anti-αVβ5. **g**) The percentage of cell spreading on FHL-1 in the presence of each inhibiting antibody was calculated. The inhibitory effects of both anti-β1 and anti-α5 on primary RPE cell spreading on FHL-1 were shown to be dose dependent in (**h**) and (**i**) respectively. Images in **a-f** are representative of 3 independent experiments. Data in **g** represent n=3 ± s.e.m. Statistical analysis was performed by Student T test, where *=*P*<0.05 and ****=*P*<0.0001. Data in **h-i** are n=3 ± s.e.m. Scale bar represents 100μm.

In additional competition experiments, a cyclic GRGDS peptide decreased the percentage of spread RPE cells on FHL-1 compared to a reverse-sequence control peptide (SDGRG) (Figure 4a-b). To confirm the role of the RGD domain in FHL-1 as an integrin binding site an FHL-1 RGD ‘null’ protein was made, where the aspartate residue in the RGD sequence was mutated to an alanine residue (making an RGA sequence, see Figure 4a). The FHL-1-RGD null protein retained its functional co-factor activity for the complement factor I (FI) mediated breakdown of C3b into iC3b (see Supplementary Figure 3), but lost all capacity to support primary RPE cell spreading (Figure 4c-g). This lack of interaction was not due to the presence of any endotoxin contamination within the recombinant protein preparations as the measured endotoxin levels for FHL-1 and FHL-1 RGD null were 0.02 ng/ml and 0.03 ng/ml, respectively.

**Figure 4.**
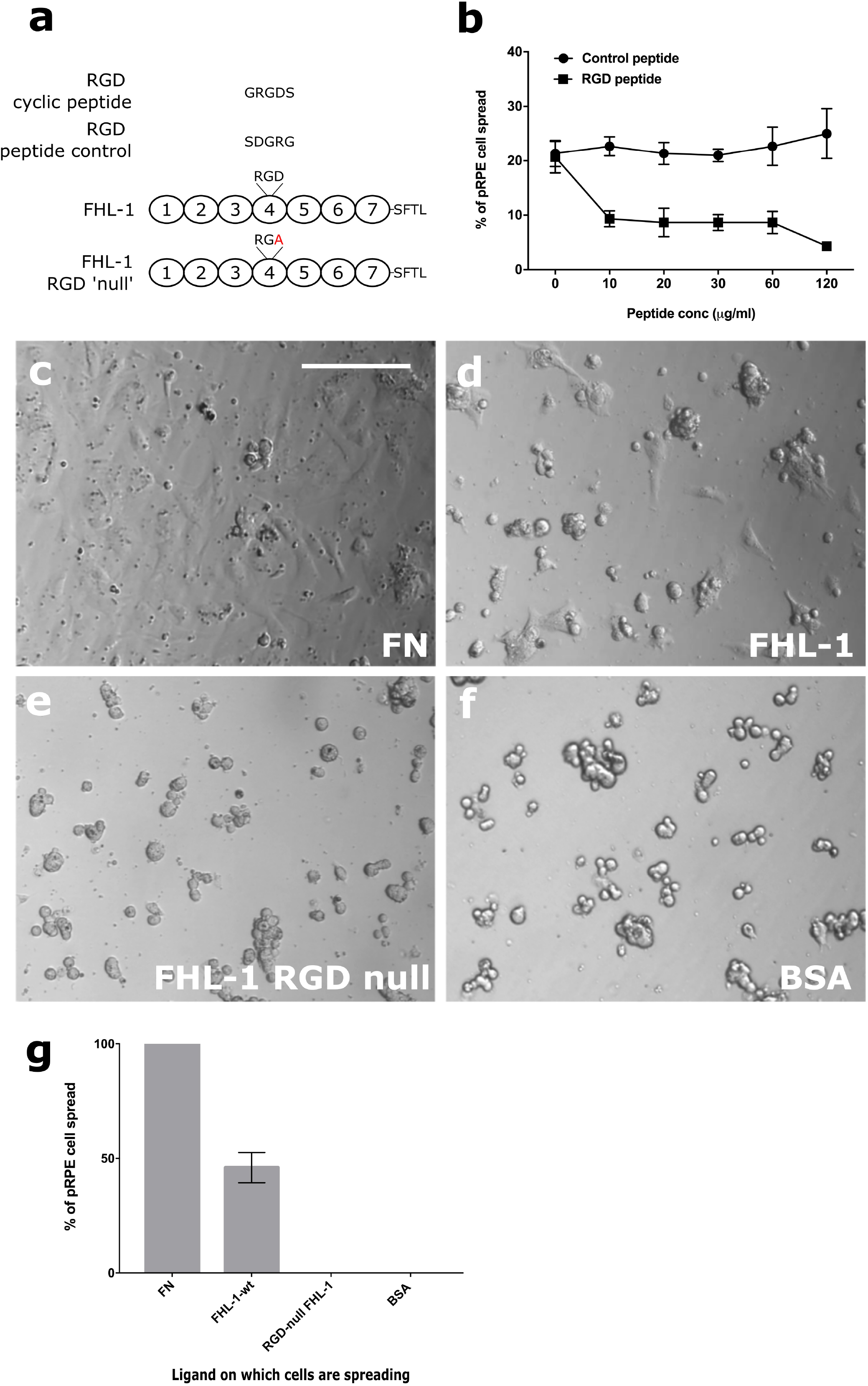
RPE cell α5β1 integrin recognises the RGD binding motif in FHL-1. **a**) Schematic showing the design of both the RGD cyclic peptide and the scrambled peptide control, as well as the location of the aspartic acid mutation to an alanine residue for the creation of the RGD null FHL-1 protein. **b**) The spreading of primary RPE cells to immobilised FHL-1 was inhibited with increasing concentrations of the cyclic RGD peptide: no inhibition was observed under the same conditions with the control scrambled peptide. Data comprises n=3 ± s.e.m. The binding of the α5β1 integrin to the RGD binding domain in FHL-1 is tested by incubation of primary RPE cells with, **a**) fibronectin, **b**) FHL-1, **c**) FHL-1 RGD null, or **d**) BSA. **e**) Percentage of spread cells were calculated and compared to the positive control, fibronectin. Images in **a**-**d** are representative of 3 independent experiments. Data in **e** represent n=3 ± s.e.m. Scale bar represents 100μm.

### The hTERT-RPE1 cell line mimics primary RPE cell behaviour

Next, we investigated the effects on gene transcription of the integrin/FHL-1 interaction, but surmised that this would be challenging using primary cells due to the naturally occurring donor-to-donor variability. Therefore, we tested the suitability of the RPE cell line hTERT-RPE1 for use in further experiments. Cell spreading assays were repeated as before and approximately 70% cell spreading was demonstrated on immobilised FHL-1 with hTERT-RPE cells when compared to FN (Figure 5). No cell spreading was observed on the FHL-1 RGD null mutant control, confirming that the interaction was RGD-binding integrin mediated.

**Figure 5.**
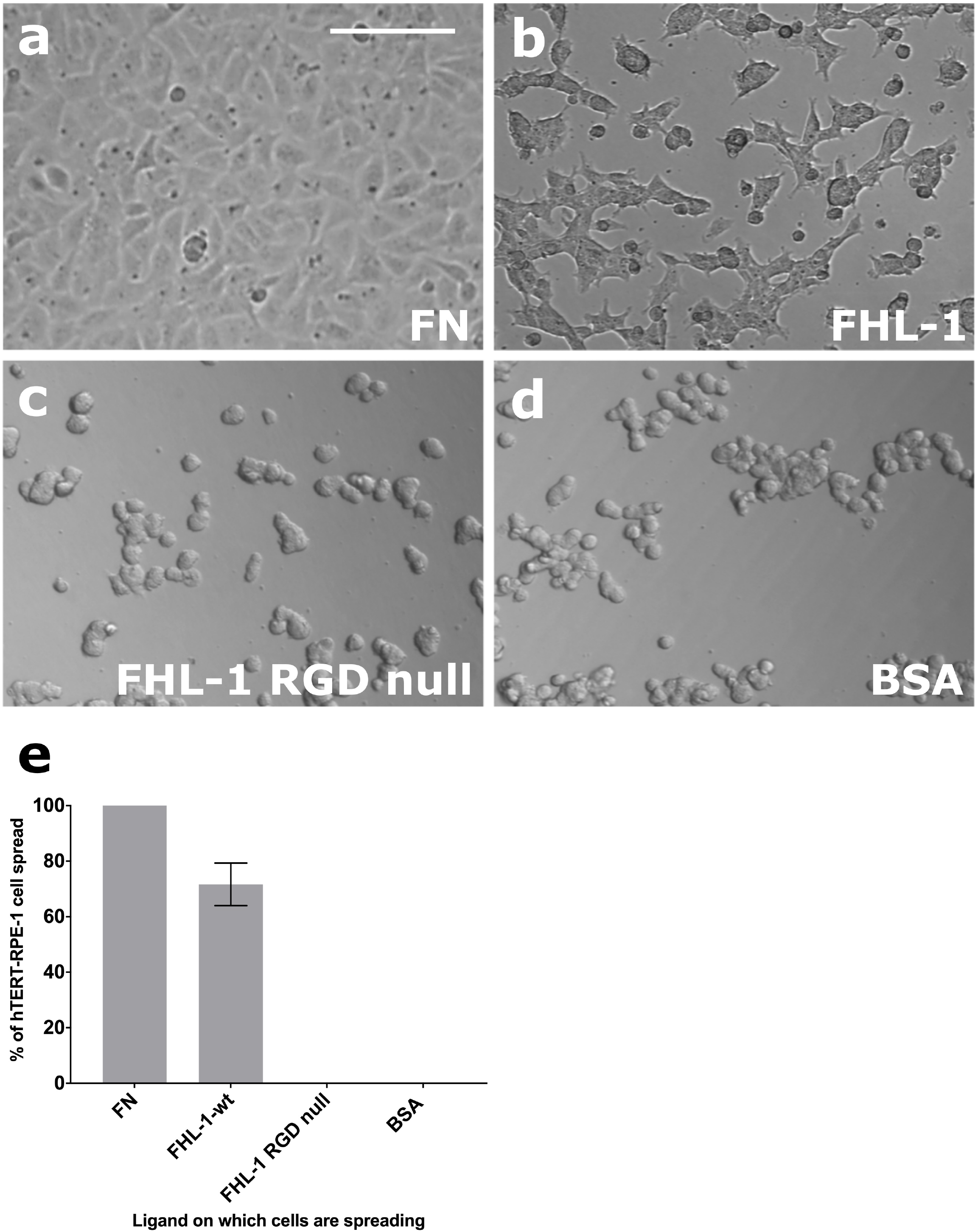
hTERT-RPE1 cells mirror the FHL-1 binding of primary RPE cells. The hTERT-RPE cell line was tested for FHL-1 binding by addition to different matrices for three hours, including **a**) FN; **b**) FHL-1; **c**) FHL-1 RGD null; and **d**) BSA. **e**) the number of spread cells were counted in four different visual fields for each condition and each condition were measured in triplicate. The percentage of spread cells was calculated and compared to the positive control, FN. Images in **a**-**d** are representative of 3 independent experiments. Data in **e** represent n=3 ± s.e.m. Scale bar represents 100μm.

### Substrate-dependent alteration of gene expression in hTERT-RPE1 cells

In order to investigate the effects of different substrates on gene expression RNA-seq transcriptome analysis was carried out on hTERT-RPE1 cells grown for 24h on three different substrates; FHL-1, FN and LA. Data were analysed using Ingenuity Pathway Analysis (IPA; Qiagen) software comparing differences between substrate in a core analysis with a false discovery rate (FDR) of 0.1. A further analysis was carried out using DAVID (https://david.ncifcrf.gov/): see Supplementary Tables 1-3.

For functional annotation clustering in DAVID analysis, gene lists were constructed comparing differential gene expression in cells between the different substrates: FHL-1 v FN (201 differentially expressed genes), FHL-1v LA (139 genes) and LA v FN (162 genes), see Supplementary Tables 1-3. Functional annotation clustering, Supplementary Tables 4-6, revealed that the majority of changes when comparing FHL-1 and FN concerned cell cycle and DNA replication, while in the case of comparing FHL-1 and LA, the highest enrichment occurred in the immune response and cytokine activity.

IPA analysis also revealed similar differences in canonical pathways predicted to be altered when comparing between substrates (Figure 6: all significantly altered pathways of interest are found in Supplementary Tables 7-9). In the case of FHL-1 v FN, the majority of genes in the top 20 predicted pathways were found to be downregulated by FHL-1 especially those involved in the cell cycle and/or DNA repair, e.g. cell cycle control of chromosomal replication, mitotic roles of polo-like kinase, cell cycle G2/M DNA damage checkpoint regulation, telomere extension by telomerase, and role of CHK proteins in cell cycle checkpoint control. When comparing FHL-1 v LA, the majority of genes were upregulated by FHL-1 in predicted pathways, these included cell/cell interaction pathways such as axonal guidance, and integrin signalling. There were no differences observed in integrin signalling when comparing FHL-1 v FN, suggesting these two substrates regulate similar genes in that pathway. Additional pathways predicted to be altered in FHL-1 v LA included intracellular cell signalling pathways, e.g. EIF2 signalling, mTOR signalling, nuclear receptor signalling pathways (glucocorticoid and aldosterone), Rho signalling pathways (RhoA, RhoGDI and actin-based motility by Rho). The NRF2-mediated oxidative stress response was also altered, suggesting that FHL-1 may exert anti-oxidant effects. A number of these pathways were commonly regulated in FHL-1 v FN and FHL-1 v LA but not in LA v FN, and therefore are likely to be specific to FHL-1, i.e. unfolded protein response, aldosterone signalling in epithelial cells, and EIF2 signalling. Taking these data together, FHL-1 predominantly inhibits expression of genes controlling the cell cycle compared to FN and exhibits differences in cell/cell interaction mechanisms when compared to the effects of LA.

**Figure 6.**
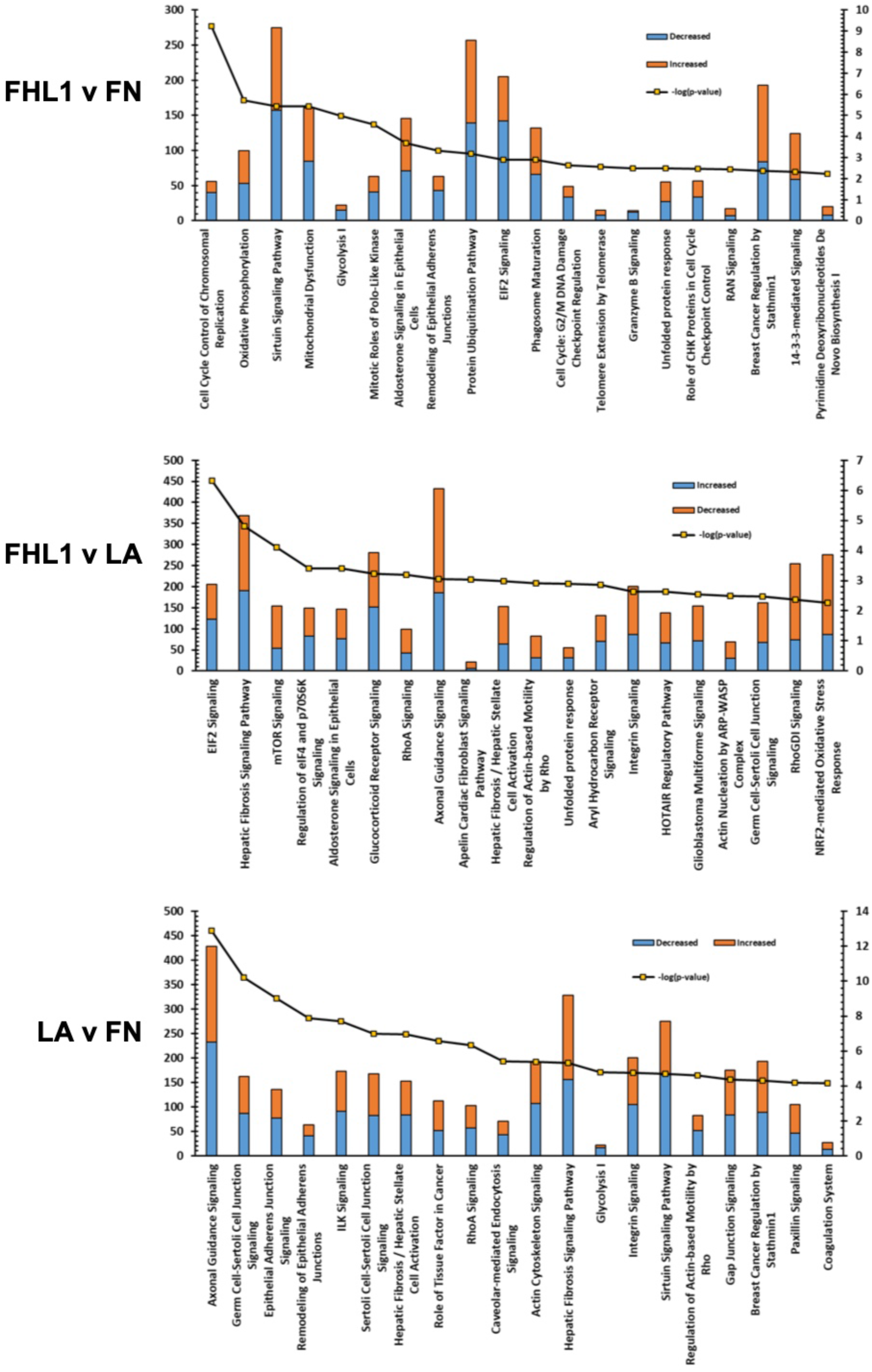
IPA analysis of differentially expressed genes in the most significantly altered canonical pathways. Bar charts indicate the most significantly different canonical pathways and the number of differentially expressed genes for each substrate comparison. Blue indicates decreased expression; orange genes indicate increased expression according to each comparison (measured on the left axis) and the line indicates significance as −log p value (measured on the right axis).

### FHL-1 protects hTERT-RPE1 cells from oxidative stress induced cell death

As FHL-1 modifies oxidative stress gene expression, and the unfolded protein response via heat shock protein gene expression (including the *HSPA6* gene), we investigated the putative protective response of FHL-1 to oxidative stress-induced cell death. To this end, cell survival assays were performed on hTERT-RPE1 cells grown in wells pre-coated with FHL-1, RGD-null, FN, LA and PBS control for 24 hours at concentrations similar to those used for the cell spreading assays. After switching to serum-free medium supplemented with B27 without antioxidants, cells were maintained with H_2_O_2_ (150μM) for a further 24h. After this, cells were stained with Hoechst 33342 and imaged using an automated scanner and counted using ImageJ. H_2_O_2_ treatment resulted in robust cell death in control cells (PBS coated wells) compared to untreated cells (Figure 7). The number of cells in untreated wells were not significantly different when comparing different substrates. However, cells grown on FHL-1 and LA were significantly protected (p<0.01 and p<0.02 respectively) against H_2_O_2_-mediated cell death compared to FN which exhibited no such protection. FHL-1 RGD-null did not show a significant protective effect. These data identify a role of FHL-1 in mediating RPE cell survival against oxidative stress similar to that conferred by LA binding.

**Figure 7.**
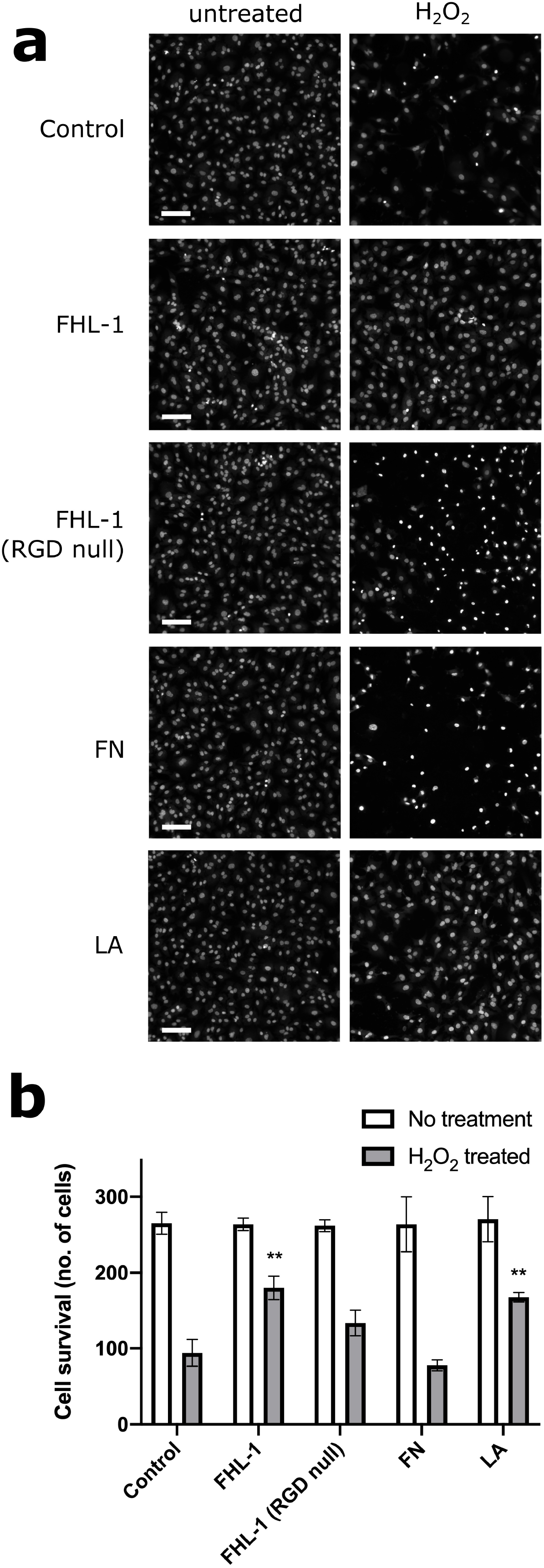
FHL-1 and LA protect hTERT-RPE1 cells from oxidative stress-induced cell death. **a**) UV Photographs of cells grown for 24h on FHL-1, FHL-1 RGD null, LA and FN coated plates subsequently stained with Hoechst 33342 after 24h in the presence or absence of H_2_O_2_. Scale bar equals 100μm. **b**) Bar chart quantifying this data demonstrates that FHL-1 and LA exert significant protective properties (p<0.01 and p<0.02, respectively) compared to FN and a partial, but non-significant protective effect with FHL-1 NULL (p>0.05)

## Discussion

Here, we describe a novel interaction between human RPE cells and a protein associated with Bruch’s membrane, the complement regulator FHL-1 (7, 18). Despite the previously known important complement regulatory functions of FHL-1, the interaction described in this study demonstrates a non-canonical role of FHL-1 in protecting RPE cells from oxidative stressed-induced cell death. The human retina is one of the most metabolically active sites within the human body and the RPE cells are particularly subject to extensive levels of oxidative stress. There is strong evidence implicating oxidative stress in the pathogenesis of AMD (32–34), and the stability of RPE cells *in vivo* is dependent on their interactions, not only with one another, but also their underlying ECM, Bruch’s membrane.

RPE cell attachment to ECM components through integrins is a well-documented phenomenon (13, 14), and increasing RPE cell integrin expression has been postulated as a method for improving the adhesion of transplanted RPE cells to a recipient’s Bruch’s membrane (35). Furthermore, integrin-mediated RPE adhesion to Bruch’s membrane confers protective effects (36, 37). Previous work investigating these interactions has focused on the main ligands of integrin receptors within Bruch’s membrane; collagen, LA, and FN (6). By using cultured primary RPE cells, isolated from human donor eyes, we demonstrate their ability to interact with immobilised FHL-1 (see Figure 2). Given that FHL-1 is identical to the first seven domains of FH (plus a unique four amino acid C-terminal tail: see Figure 1b), and both have exactly the same RGD-containing CCP4 (16, 17), it was of interest that RPE cells spread on immobilised FHL-1 but not FH. This phenomenon has been observed previously where a series of anchorage-dependent cell lines were shown to adhere to FHL-1, but not FH (27). This study, which used recombinant fragments of the FHL-1 protein, identified the RGD binding domain of FHL-1 as being essential for these interactions. Furthermore, without direct testing the authors hypothesised that integrins were responsible given the reliance on bivalent metal ions for the interactions to be successful. Interestingly, they also observed ~50% cell spreading on FHL-1 when compared to FN, where in our study we observed ~40% with cultured primary RPE cells and ~70% with the hTERT-RPE cell line (Figures 2 and 5). However, our subsequent competition of RPE cell/FHL-1 interactions with FH in the fluid phase (Supplementary Figure 2) suggests that the lack of RPE cell interaction with immobilised FH is a result of how the protein adheres to plastic in our experimental settings rather than necessarily a representation of its lack of RPE cell adhesion *in vivo*. The predominance of FHL-1, rather than FH, within Bruch’s membrane (18) may indicate that the RPE/ FHL-1 interaction is more important for oxidative stress induced gene expression, although this does not exclude a lesser involvement of FH in RPE cell resilience to oxidative stress.

Evidence of complement over-activation in the ECM of the choriocapillaris in AMD, and indeed preceding clinical manifestation of the disease, suggests complement gene variants that confer AMD-risk cause this dysregulation (38–42). In addition, focus has recently been given to the non-canonical roles of complement proteins in AMD pathogenesis. For example, the common high-risk Y402H polymorphism in FH has been associated with lipoprotein dysregulation and causing an ocular phenotype in aged mice (26). Also, the Y402H polymorphism hinders the ability of FHL-1 to bind glycosaminoglycans (GAGs), such as heparan sulphate (18, 43–45), although the presence of a secondary GAG-binding site in the full-length protein means FH itself is not affected (46). The age-associated reduction in heparan sulphate chains in Bruch’s membrane (47), coupled with the Y402H predisposition to weaker binding, means less FHL-1 will be present in Bruch’s membrane to regulate local C3b deposition and complement activation. This study suggests that the decrease in ECM bound FHL-1 would also result in less RPE cell interactions with FHL-1 and consequently increase their susceptibility to oxidative stress.

We investigated the effects of different immobilised integrin ligands on hTERT-RPE cell gene transcription using FHL-1, FN and LA substrates. FN binds to similar integrins as FHL-1 (*ITGA5:ITGB1*) while LA does not. RNAseq, Ingenuity Pathway Analysis, as well as functional annotation clustering using DAVID, uncovered altered activity in pathways that were specific to FHL-1 when compared to either FN or LA. Compared to FN (which was not protective in the cell survival assays), cells grown on FHL-1 exhibited changes in genes associated with the cell cycle, including: Cell Cycle Control of Chromosomal Replication, Mitotic Roles of Polo-Like Kinase, Cell Cycle: G2/M DNA Damage Checkpoint Regulation and Role of CHK Proteins in Cell Cycle Checkpoint Control (see Supplementary Table **7**). The majority of genes in these pathways were down-regulated, but any putative effect on cell cycle by FHL-1 is likely to be subtle, as there were no differences in cell numbers when comparing hTERT-RPE cells on FHL-1 and FN in the cell survival assays (when not treated with hydrogen peroxide) (Figure 7). Although there have been no previous studies on the role of FHL-1 on cell cycle control, mice deficient of full-length FH (also not expressing FHL-1) exhibit changes in the number of cells in the developing retina (25). These *Cfh*^-/-^ animals displayed reductions in the rate of mitosis during a critical period of retinal development directly after birth. It is interesting to note that this study also observed enlarged mitochondria, observations consistent with premature ATP decline, degeneration and/or senescence. Ingenuity Pathway Analysis also identified changes in metabolic pathways involved in ATP production such as oxidative phosphorylation and glycolysis as well as mitochondrial dysfunction and sirtuin signalling, which is responsible for aging/senescence processes within the cell (48). In relation to AMD, mitochondrial dysfunction and DNA damage occurs in the disease-associated Y402H variant of FH (49) and this is presumed to be due to the increase in formation radical oxygen species and subsequent oxidative stress.

Compared to hTERT-RPE cells grown on LA, the cells grown on FHL-1 exhibited altered expression in a number of pathways that were different to those observed when comparisons were made to FN. Interestingly, integrin signalling was one such pathway, and a component of that pathway, ITGA5 was found to be a significantly differentially expressed. FHL-1 and FN both bind to ITGA5:ITGB1. Therefore, it was interesting to observe when comparing gene expression changes in cells grown in LA with those grown on FN, the integrin pathway signalling was found to be an affected pathway, highlighting that FHL-1 and FN share common mechanisms in regulating integrin signalling. FHL-1 v LA-specific changes included mTOR Signalling, a pathway important in maintaining retinal homeostasis in RPE cells by regulating lysosomal phagocytosis and autophagy (50), and NRF2-mediated Oxidative Stress Response that may confer protection to oxidative insults (51). Studies have previously linked abnormal autophagy with reduced NRF2 signalling in animal models of AMD (52).

HSPA6, which is an inducible form of HSP70, was found to be the highest upregulated gene when cells were grown on immobilised FHL-1 when compared to either FN or LA. HSP6A which is only partially conserved in the mammalian lineage and has no homologs in rodents has been reported to be induced in an *in vitro* model of photocoagulation in the ARPE-19 cell line (53). HSPA6 and another inducible HSP70 gene, HSP1A, which was also demonstrated to have increased expression in cells grown on FHL-1, work in tandem to protect cells against proteotoxic insults and heat shock-mediated cell death in various cell lines (54, 55). HSP70 proteins are also known to interact with mineralocorticoid receptor (MR), a high affinity ligand of aldosterone, and keep it in a basal state (56). In this non-activated state, MR is predominantly cytoplasmic and part of a large heteromeric complex interacting with a number of proteins including HSPs. Upon ligand binding, a conformational change occurs that leads to the dissociation of the complex and subsequently the MR translocates to the nucleus binding to DNA leading to the regulation of gene transcription. Aldosterone binding and activation of MR signalling pathways are associated with increased levels in oxidative stress in vascular inflammation (57) and this is relevant to certain retinal disorders. Use of the MR antagonist spironolactone, reduces CNV activity in patients with refractory neovascular AMD in a VEGF-independent manner (58). Therefore, an increase in the level of heat shock proteins (and thus increased sequestration of MR to the cytoplasm) is one potential mechanism by which FHL-1 may illicit a protective response to RPE cells from oxidative insult.

These findings point to a putative role of FHL-1 in mediating a response to oxidative stress. Therefore, we tested the effect conferred by FHL-1 with RPE cell survival in a hydrogen peroxide-induced oxidative stress model. While immobilised FHL-1 exhibited protective effects similar to that observed with immobilised LA, cells grown on FN did not survive to the same extent (Figure 7). This indicates that any mechanism of FHL-1 mediated protection occurs independently of any interaction with the integrin signalling pathway shared with FN. The protective effect is not due to changes in cell cycle, as the number of cells that grew on immobilised FHL-1 (and untreated with hydrogen peroxide) were similar to those of untreated cells grown directly on plastic. No previous studies have demonstrated that FHL-1 can confer a protective effect against oxidative stress; however, a recent study has shown that full length FH, when supplemented to the cell culture medium can protect ARPE-19 and human iPSC-derived RPE cells from oxidative insult as a result of 4-HNE treatment (59). Furthermore, Borras *et al*. found that FH inhibits caspase-induced apoptosis and protects RPE tight junctions from oxidative stress-induced disruption (59). However, the Borras *et al*. study did not identify how the full-length protein was interacting with the RPE cells to confer these effects as recombinant truncated fragments of FH all failed to replicate the results seen with the full-length protein.

Our study has been the first to demonstrate a role of FHL-1 in providing RPE cell resistance to oxidative stress, and highlights the need to fully understand how RPE cells interact with their underlying ECM. Here we show that FHL-1 alters the gene transcription of RPE cells by binding to their α5β1 integrin receptor. This adds a novel function to the repertoire of FHL-1 by mediating both a complement response as a co-factor for FI in this micro-environment, but also confer a protective effect against oxidative insults on the RPE cells. This novel finding will help in our understanding of RPE cell behaviour *in vivo*, and highlights the need to consider the endogenous RPE cell/ECM interactions when designing therapeutic interventions.

## Abbreviations

AMD: Age-related macular degeneration
CCP: Complement control protein
DMEM: Dulbecco’s modified eagle’s medium
ECM: Extracellular matrix
FCS: Foetal calf serum
FDR: False discovery rate
FH: Factor H
FHL-1: Factor H-like protein 1
FHR-4: Factor H-related protein 4
FI: Factor I
FN: Fibronectin
GAGs: Glycosaminoglycans
LA: Laminin
LAL: Limulus amebocyte lysate
MR: Mineralocorticoid receptor
NSR: Neurosensory retina
NGS: Normal goat serum
PFA: Paraformaldehyde
RPE: retinal pigment epithelium

## Acknowledgements

We thank Leo Zeef and Andy Hayes of the Bioinformatics and Genomic Technologies Core Facilities at the University of Manchester for providing support with regard to RNA-seq. This work was funded by a Fight for Sight research grant (1852/53) and an MRC Career development Fellowship (MR/K024418/1). SJC is funded by the Helmut Ecker Foundation, Germany.

## Conflicts of Interest Statement

PNB, and SJC are inventors of patent applications that describe the use of complement inhibitors for therapeutic purposes, and are co-founders and Directors of Complement Therapeutics, which is developing complement inhibitors for therapeutic purposes. RC, NB, RS, ES, VT, MSU, SM, JAA and MJH all declare that they have no conflicts of interest.

## Author Contributions

RC, NB, PNB and SJC designed research; RC, NB, RS and SJC performed research; ES, VT, MSU, SM, JAA and MJH contributed new reagents or analytical tools; RC, NB, PNB and SJC analysed data; RC, NB, PNB and SJC wrote the paper. SJC co-ordinated the project and all authors contributed to the final version of the manuscript text.

## Notes

### Competing Interest Statement

Paul Bishop, and Simon Clark are inventors of patent applications that describe the use of complement inhibitors for therapeutic purposes, and are co-founders and Directors of Complement Therapeutics, which is developing complement inhibitors for therapeutic purposes. All other authors declare that they have no conflicts of interest.

