## Supplemental text for "FHL-1 interacts with human RPE cells through the α5β1 integrin and confers protection against oxidative stress"

**Supplementary Table 1.** List of differentially expressed genes in hTERT-RPE cells cultured on FHL-1 v FN.

**Supplementary Table 2.** List of differentially expressed genes in hTERT-RPE cells cultured on FHL-1 v LA.

**Supplementary Table 3.** List of differentially expressed genes in hTERT-RPE cells cultured on LA v FN.

**Supplementary Table 4.** Functional annotation clustering for gene expression changes in hTERT-RPE cells cultured on FHL-1 vs FN.

**Supplementary Table 5.** Functional annotation clustering for gene expression changes in hTERT-RPE cells cultured on FHL-1 vs LA.

**Supplementary Table 6.** Functional annotation clustering for gene expression changes in hTERT-RPE cells cultured on LA vs FN.

**Suppplementary Table 7.** Significantly altered pathways and genes found to be altered using IPA analysis, FHL-1 v FN

**Suppplementary Table 8.** Significantly altered pathways and genes found to be altered using IPA analysis, FHL-1 v LA

**Suppplementary Table 9.** Significantly altered pathways and genes found to be altered using IPA analysis, LA v FN

**Supplementary Figure 1. Immunofluorescence analysis of primary RPE cell cultures.** Cultured primary RPE cells from human donor eyes were tested for the expression of tight junctions and RPE cell markers **a**) DAPI alone **b**) ZO-1, **c**) RPE65 and **d**) Bestrophin-1. Positive cell staining for each respective marker is shown in green, and DAPI staining of cell nuclei is shown in blue. Images are representative of 3 individual experiments.

**Supplementary Figure 2. Fluid phase inhibition of primary RPE cell spreading on immobilised FHL-1 with FN, FH and FHL-1 proteins.** The spreading of primary RPE cells on FHL-1 was inhibited by the addition of fluid phase FHL-1 and FN, but not a recombinant protein comprising solely CCP6-7 of FH. Data is n=3 ± s.e.m. Statistical analysis was performed by Student T test, where **=*P*<0.01 and ***=*P*<0.001.

**Supplementary Figure 3. C3b breakdown assay to assess the functional capacity of FHL-1 RGD null protein.** Both wild-type FHL-1 and RGD-null FHL-1 were incubated with C3b and FI (see materials and methods). After incubation, samples were run on a reducing gradient gel for the separation of the protein bands. RGD-null FHL-1 (RGD to RGA) did not alter the ability of FHL-1 to contribute FI-mediated breakdown of the C3b α-chain by FI, leading to the appearance of bands at 68 and 43 kDa, which is a measure of cofactor activity. There is no difference in the functional ability of the wild-type or RGD-null FHL-1 forms. This image is representative of three independent experiments.

**Supplementary Figure 1**

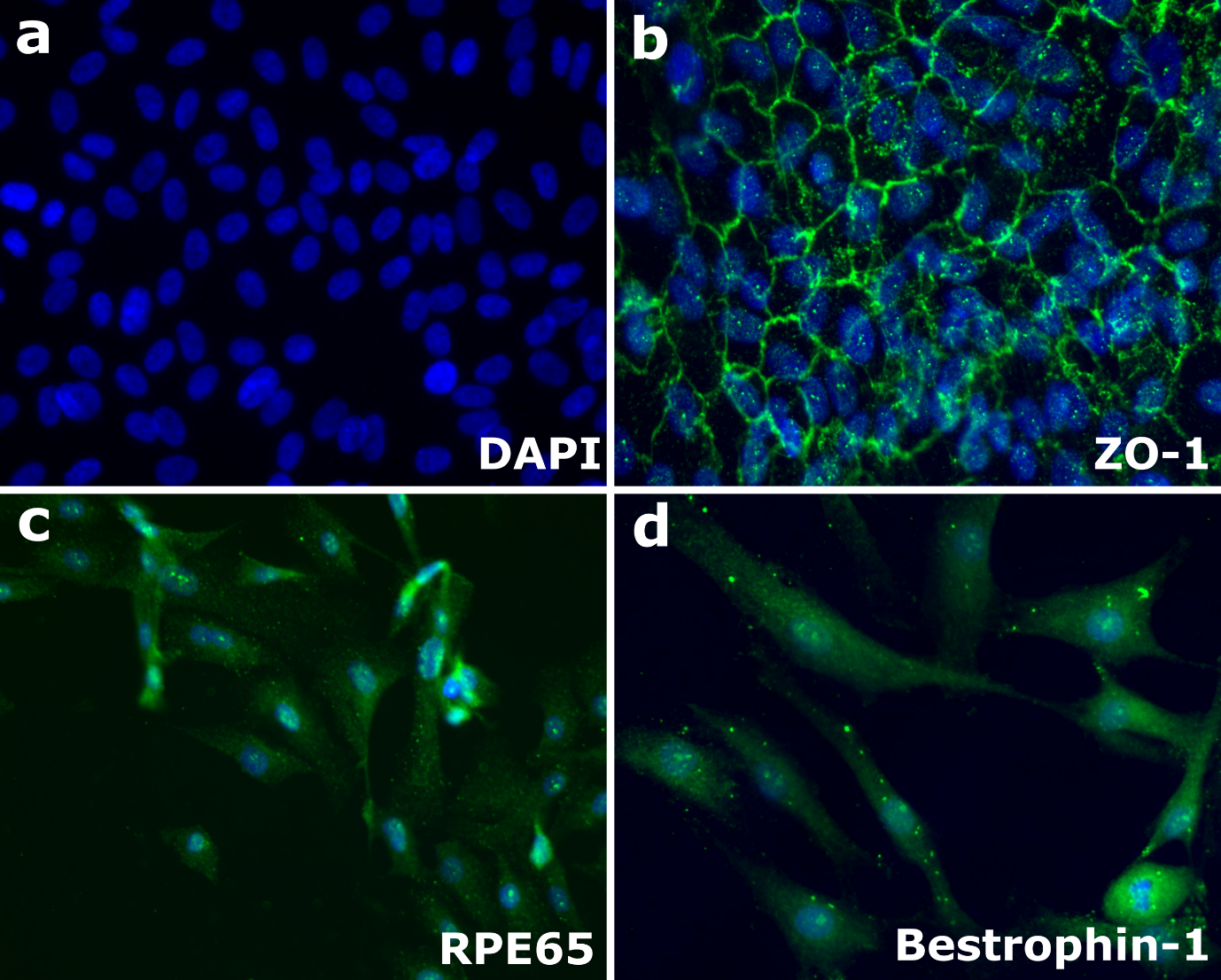

**Supplementary Figure 2**

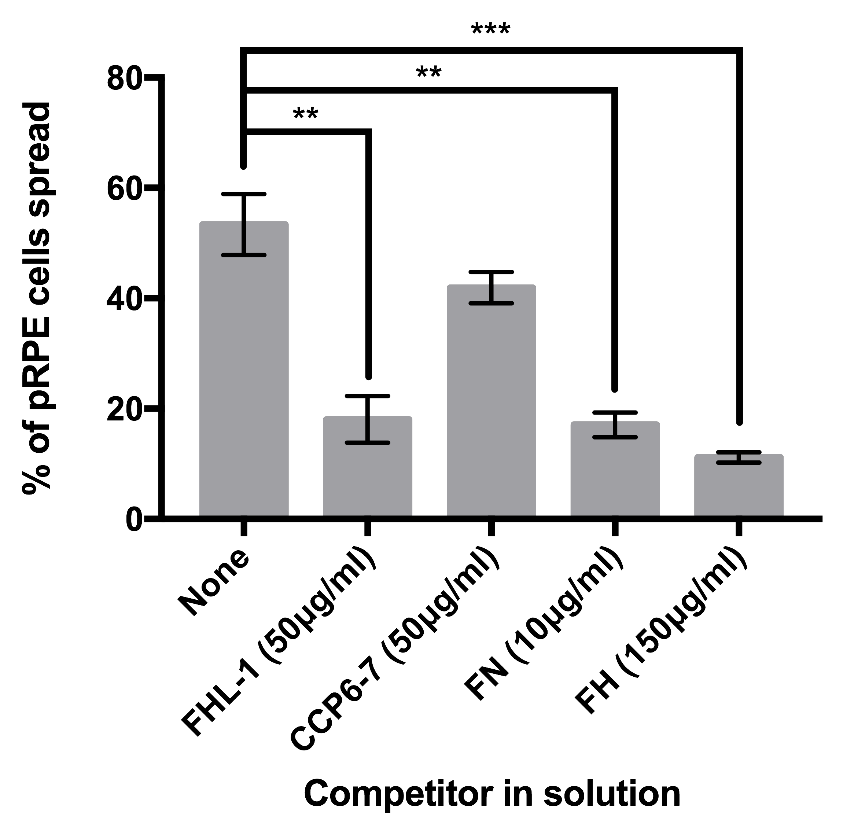

**Supplementary Figure 3**

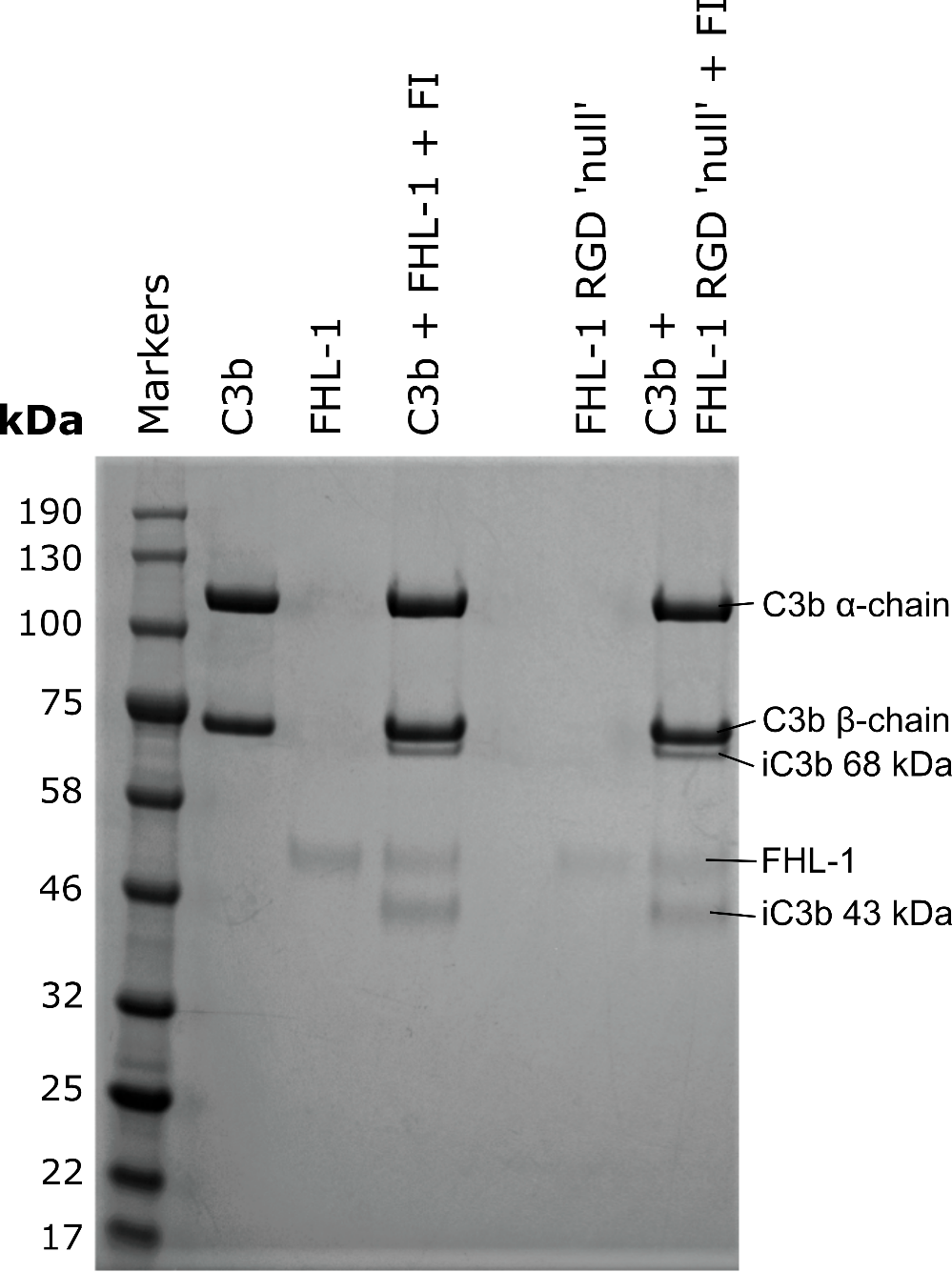
